# Non-coding RNAs Associated with Prader-Willi Syndrome Regulate Transcription of Neurodevelopmental Genes in Human Induced Pluripotent Stem Cells

**DOI:** 10.1101/2021.05.11.443612

**Authors:** Monika Sledziowska, Matt Jones, Ruba Al Maghrabi, Daniel Hebenstreit, Paloma Garcia, Pawel Grzechnik

## Abstract

Mutations and aberrant gene expression during cellular differentiation lead to neurodevelopmental disorders such as Prader-Willi syndrome (PWS) which results from the deletion of an imprinted locus on chromosome 15. We analysed chromatin-associated RNA in human induced pluripotent cells (iPSCs) upon depletion of hybrid small nucleolar long non-coding RNAs (sno-lncRNAs) and 5’ snoRNA capped and polyadenylated long non-coding RNAs (SPA-lncRNAs) transcribed from the locus deleted in PWS. We found that rapid ablation of these lncRNAs affects transcription of specific gene classes. Downregulated genes contribute to neurodevelopment and neuronal maintenance while genes that are upregulated are predominantly involved in the negative regulation of cellular metabolism and apoptotic processes. Our data revealed the importance of SPA-lncRNAs and sno-lncRNAs in controlling gene expression in iPSCs and provided a platform for synthetic experimental approaches in PWS studies. We conclude that ncRNAs transcribed from the PWS locus are critical regulators of a transcriptional signature important for neuronal differentiation and development.

## INTRODUCTION

Prader-Willi syndrome (PWS) is a genetic neurodevelopmental disorder characterised by hypotonia in infancy, developmental delay, cognitive disability, behavioural problems and hyperphagia often leading to life-threatening obesity (Angulo et al., 2015). The cause of PWS is the lack of expression of genes from the paternally inherited locus q11-q13 on chromosome 15. This can occur as a result of a paternal deletion in the 15q11-q13 region (70% of cases), maternal uniparental disomy (20-30% of cases) or imprinting defect (1% of cases) (Cheon, 2016). *Post-mortem* analysis of hypothalamic tissue of patients with PWS revealed upregulation of genes signalling hunger and downregulation of genes which regulate feeding (Bochukova et al., 2018). Moreover, genes involved in neurogenesis, neurotransmitter release and synaptic plasticity were downregulated in those patients, while microglial genes associated with inflammatory responses were excessively expressed (Bochukova et al., 2018). The molecular processes resulting in this misregulation of gene expression are yet to be determined.

The PWS locus encodes the *SNURF-SNRPN, NDN, MKRN3, NAPAP1*, and *MAGEL2* genes. *SNURF-SNRPN* 3’ untranslated region extends into the non-coding region called *SNGH14*. The introns of *SNGH14* contain multiple clusters of box C/D small nucleolar RNAs (snoRNAs) (Qi et al., 2017). Both *SNURF-SNRPN* and *SNHG14* share the same promoter and exons. The minimal deletion associated with PWS spans 118kb in *SNHG14* and encompasses twenty-nine copies of snoRNA SNORD116 and a single snoRNA, SNORD119A (Bieth et al., 2015). Box C/D snoRNAs are short (60-300 nucleotides) ncRNAs that form ribonucleoprotein complexes and mediate ribose 2’-O-methylation of predominantly ribosomal RNA (rRNA) and small nuclear RNAs (snRNAs) (Kufel and Grzechnik, 2019). These snoRNAs contain guiding sequences which are complementary to their RNA targets. However, the majority of snoRNAs in humans do not possess a clear affinity to any cellular RNA sequences and thus are called orphan snoRNAs. These snoRNAs may act on multiple RNA targets or play other undetermined roles in the cell. Similarly, snoRNAs encoded from the 15q11-q13 locus are also orphan snoRNAs and their functions remain largely unknown.

Recent studies showed that some snoRNAs from the locus missing in PWS can form two types of hybrid long non-coding RNAs (lncRNA): five small nucleolar RNA related long non-coding RNAs (sno-lncRNAs) and two 5’ snoRNA capped and polyadenylated lncRNA (SPA-lncRNAs) (Figure 1A) (Wu et al., 2016; Yin et al., 2012). Both sno-lncRNAs and SPA-lncRNAs are by-products of *SNURF-SNRPN-SNGH14* processing (Wu et al., 2016; Yin et al., 2012). Sno-lncRNAs are formed by two snoRNA embedded into the same intron which when spliced out is degraded by 3’-5’ and 5’-3’ exonucleases. Intron degradation continues until it is blocked by the snoRNA that defines the 3’ and 5’ ends of sno-lncRNAs. Thus, each of the five sno-lncRNAs consists of an intervening sequence flanked by two snoRNAs at each end, all arising from the same intron of the *SNHG14* ncRNA. The 5’ ends of the two SPA-lncRNAs in the PWS locus coincide with *SNORD107* (*SPA1-lncRNA*) and *SNORD109A* (*SPA2-lncRNA*), while the 3’ end is polyadenylated. SPA-lncRNA formation is associated with the degradation of RNA downstream of poly(A) sites (PAS). Following endonucleolytic cleavage, the RNA is degraded by the 5’-3’ exonuclease until it reaches a snoRNA sequence. This generates the 5’ end of SPA-lncRNAs and allows for Polymerase II to continue elongation to another PAS in the sequence that defines the 3’ end of SPA-lncRNAs. The intervening regions of sno- and SPA-lncRNAs were shown to sequester the RNA processing and splicing factors TDP43, RBFOX2, hnRNP M and hence affect alternative splicing (Wu et al., 2016).

**Figure 1.**
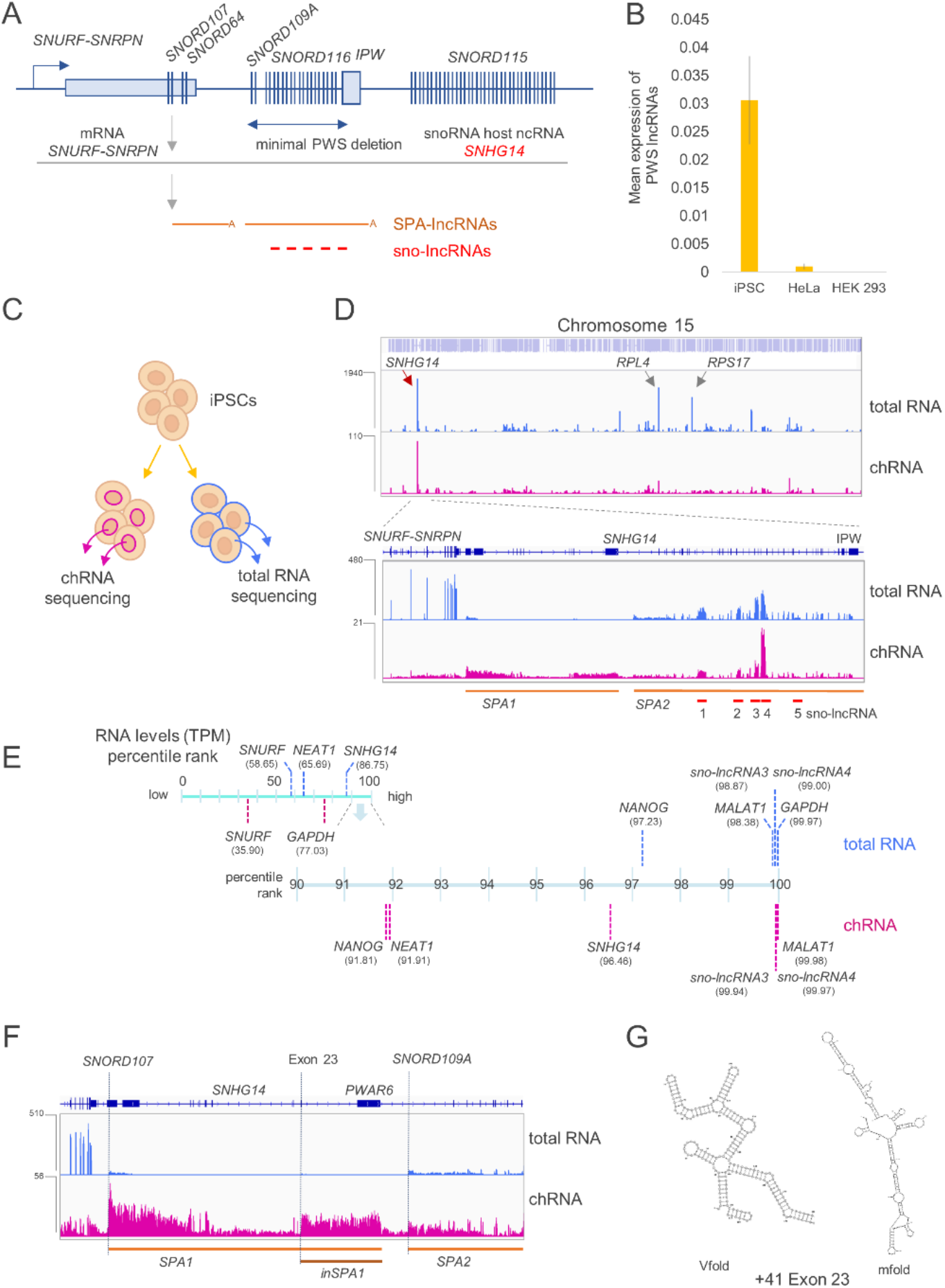
Accumulation of PWS-related ncRNAs on the chromatin in iPSCs. (A) Diagram showing organization of lncRNAs transcribed downstream of *SNURF-SNRPN* gene. *SNORD* – snoRNA genes, *IPW* – imprinted gene in the Prader-Willi syndrome region. (B) Accumulated expression of PWS lncRNAs in IPSCs, HEK293T and HeLa cells relative to the house-keeping gene *GAPDH*. The average of three biological experiments is shown, error bars indicate standard error. (C) The experimental approach used in the study. RNA was isolated either from the whole cell (total RNA) or from the insoluble nuclear fraction (chromatin-associated RNA). (D) Distribution of total RNA (blue) and chromatin-associated RNA (purple) on chromosome 15. Locations of SPA-lncRNAs and sno-lncRNAs are shown below the track; chRNA-seq analysis. (E) The abundance of RNAs in iPSCs in total RNA and chromatin-associated RNA fractions are shown for selected genes as a percentile rank. (F) Distribution of chRNA-seq reads indicating the position of putative additional lncRNA (*inSPA1*). (G) Secondary RNA structure of exon 23 revealed by Vfold and mfold predictions. chRNA-seq tracks shows counts ×10^6^; chRNA – chromatin-associated RNA; TPM – transcript per million.

Previous studies of PWS in cellular models focused primarily on changes in total RNA (Bochukova et al., 2018; McCann and Baserga, 2012; Zahova et al., 2021), which reflects the levels of cytoplasmic steady-state RNA. We examined the possibility that the ablation of sno-lncRNAs and SPA-lncRNAs from the 15q11-q13 locus affects the nascent transcriptome. Since this is usually not detectable in total RNA, which depends largely on transcript stability, we examined chromatin-associated RNA (chRNA) that reflects active nascent transcription across the genome. Acute depletion of sno- and SPA-lncRNA in human induced pluripotent stem cells (iPSCs) revealed their role in the regulation of transcription of neuronal genes. These observations provide an insight into a possible molecular mechanism by which deletion in the 15q11-q13 region affects neuronal differentiation and thus gives rise to the cognitive and behavioural symptoms occurring in PWS.

## RESULTS

### SPA-lncRNAs and sno-lncRNAs accumulate at high levels in iPSCs

SPA- and sno-lncRNAs were initially described in human embryonic H9 line and teratocarcinoma PA1 cells (Wu et al., 2016; Yin et al., 2012). Since PWS is a neurodevelopmental disease, we chose human induced pluripotent cells (iPSCs) as our model to investigate the impact of PWS-related ncRNAs on gene expression in undifferentiated cells. We employed RT-qPCR analysis to determine if SPA- and sno-lncRNAs were expressed in the iPSC (CREM003i-BU3C2) line reprogrammed from a blood sample of a 40 year old human male (Park et al., 2017) and HEK293T and HeLa cell lines derived from human embryonic kidney and cervical cancer cells, respectively. While SPA- and sno-lncRNAs were abundantly expressed in iPSCs, they were barely detectable in either HEK293T or HeLa cells (Figure 1B). This is consistent with previous report indicating that these RNAs are abundant in stem cells but not in non-pluripotent cells (Wu et al., 2016; Yin et al., 2012). Given with this observation, we confirmed iPSCs as a model suitable for PWS research.

SPA- and sno-lncRNAs are localised close to their transcription sites (Wu et al., 2016; Yin et al., 2012). Thus, we tested if these ncRNAs were bound to the chromatin in iPSCs by performing chromatin-associated RNA sequencing (chRNA-seq). We first extracted nuclei, then separated the soluble fraction and the insoluble chromatin pellet, which retains tightly associated transcription factors and chromatin-associated RNAs, including newly synthesised and nascent RNA (Figure 1C). In parallel, from intact cells, we isolated total RNA, which is dominated by steady-state, cytoplasmic RNA. Both fractions were prepared in biological duplicates and sequenced on the Illumina platform. The fractionation procedure in iPSCs was successful, the chromatin RNA fraction showed a clear increase in intronic to exonic reads ratio, indicative of the efficient removal of steady-state RNA (Figure S1A). In the total RNA fraction, we found that RNAs from *SNHG14* locus, encompassing the SPA- and sno-lncRNAs, were one of the most abundant transcripts on chromosome 15, followed by two mRNAs encoding ribosomal proteins RPL4 and RPS17. In the chromatin fraction, transcripts from *SNHG14* were the dominant RNAs from chromosome 15 (Figure 1D). In particular, *sno-lncRNA3* and *sno-lncRNA4* were the most abundant in both total, and chromatin-associated RNA. This was in contrast to data from H9 cells where SPA-lncRNAs were one the most highly expressed ncRNAs from the PWS locus only second to *SNURF-SNRPN* mRNA (Wu et al., 2016).

This observation prompted us to test the overall cellular abundance of PWS ncRNAs in iPSCs. We calculated transcript per million (TPM) values for total RNA and chromatin-associated RNA samples and ranked transcripts by their expression level (Figure 1E). In total RNA, containing mostly cytoplasmic RNAs, *SNHG14* was within 14% of the top expressed genes (ranked on 86^th^ percentile), higher than ubiquitously transcribed lncRNA *NEAT1* (65^th^ percentile). *SNURF-SNRPN* mRNA was ranked on 58^th^ percentile. One of the most abundant RNAs in the total RNA fraction were *MALAT1* and *GAPDH* (ranked on 98^th^ and 99^th^ percentile, respectively), confirming the accuracy of our analysis. In the chromatin-associated fraction *SNHG14* transcripts ascended to the top 4% (96^th^ percentile) most abundant RNAs and were ranked higher than pluripotency factor *NANOG* and *NEAT1* (both ranked on 91^st^ percentile) (Figure 1E). Similar to the total RNA fraction, *SNURF-SNRPN* chromatin-associated mRNA was ranked much lower than *SNHG14* RNA. Discrepancies in mRNA levels between the two fractions, for example for *GAPDH* (99^th^ and 65^th^ in total and chromatin-associated RNA, respectively) reflect the fact that mRNA levels in the cell are maintained not only by RNA synthesis but also by RNA stability (Singh et al., 2019). When we analysed *sno-lncRNA3* and *sno-lncRNA4* only, these RNAs were ranked in top 2% and 1% of expressed transcripts in total and chromatin-associated RNA respectively, as seen for *MALAT1*. Such high levels indicate that RNAs transcribed from *SNHG14* may play essential roles in iPSCs.

Our analysis of RNA fractions in iPSCs revealed that almost all PWS ncRNAs were clearly detected in both the total and chromatin-associated RNA fractions, with the exception of *SPA1-lncRNA*, which was less abundant in the total RNA fraction than other transcripts (Figure 1D). Moreover, we detected increased reads spanning from position +41 of exon 23 to the *PWAR6* element of *SNHG14*, within the boundaries of the *SPA1-lncRNA* (Figure 1F). This suggests the presence of a previously unannotated ncRNA, which we term ‘inside-of-SPA1-lncRNA’ (*inSPA1*), that may arise from 5’-3’ degradation of *SPA1-lncRNA* if the exonucleases are blocked by RNA structures further downstream. Computational prediction using Vfold and mfold software revealed that the 5’ end of *inSPA1-lncRNA*, containing exon 23 and its 41 upstream nucleotides, indeed folds into a number of stem-loops that may be able to block exonucleolytic trimming (Figure 1F).

### Antisense oligonucleotides-dependent depletion of PWS transcripts

Fast depletion approaches provide an opportunity to investigate the direct effects of how the reduced RNAs act in cellular pathways. To determine the effect of an acute ablation of SPA- and sno-lncRNAs on the transcriptome, we employed antisense oligonucleotides (AOS) GapmeRs (Qiagen), which targeted lncRNAs and triggered RNA cleavage by endogenous RNaseH and the subsequent degradation by exoribonucleases. We designed a panel of GapmeRs against the individual PWS ncRNAs (Figure 2A) targeting their intervening sequences located in-between terminal snoRNAs or poly(A) tail. To compare the effect of the different types of ncRNAs on transcription we used three sets of GapmeRs against: 1) all seven sno/SPA-lncRNAs, 2) five sno-lncRNAs, and 3) two SPA-lncRNAs. These were compared with a negative control, where iPSCs were treated with an equivalent amount of non-specific GapmeRs. Since some of *SNHG14* ncRNAs overlap, the GapmeR against *SPA2-lncRNA* was designed not to affect any sequences contained within sno-lncRNAs. However, all GapmeRs against sno-lncRNAs also targeted *SPA2-lncRNA*.

**Figure 2.**
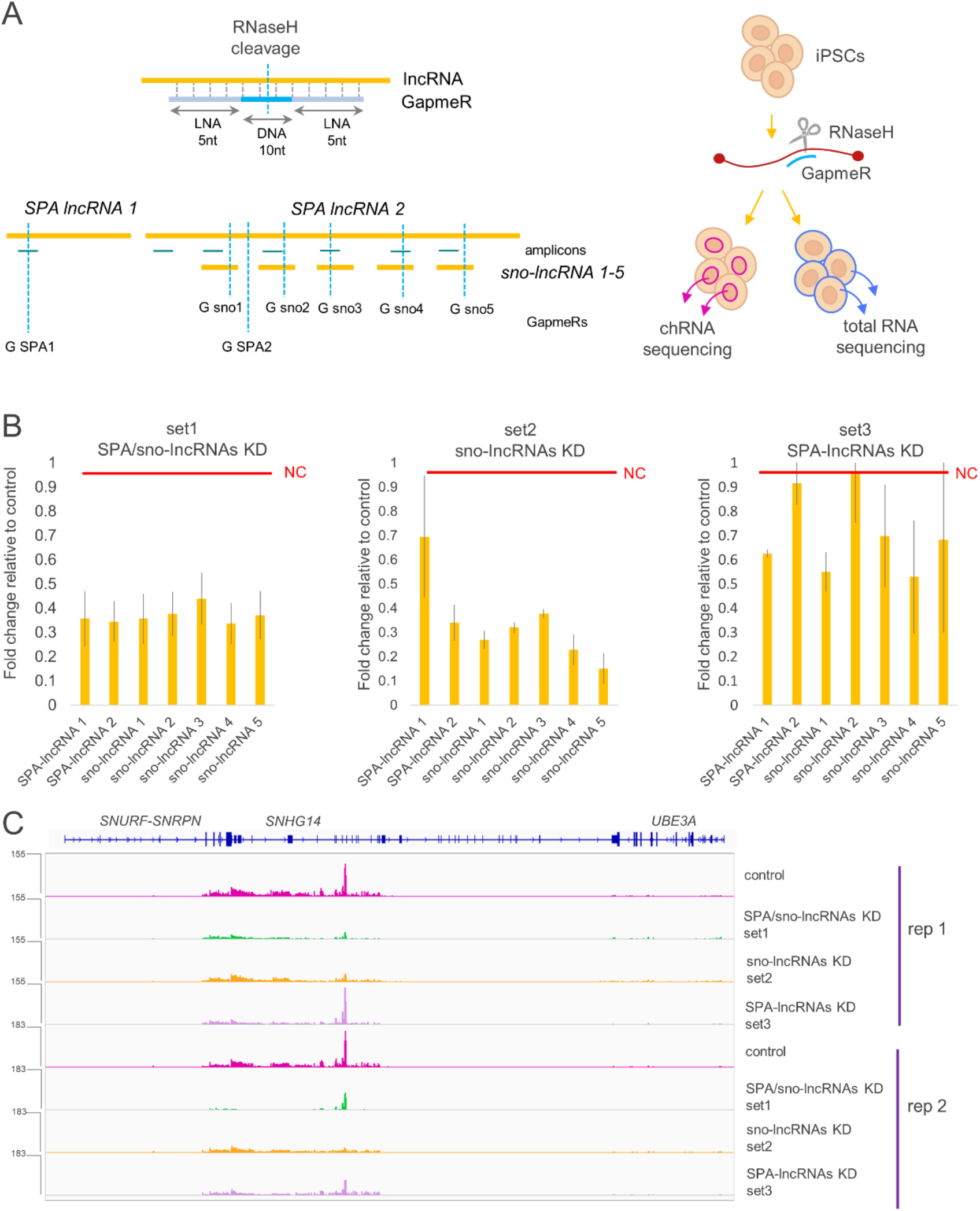
Antisense oligonucleotides mediate efficient knockdown of SPA- and sno-lncRNAs. (A) The locations of GapmeRs (vertical blue lines denoted G) targeting SPA- and sno-lncRNAs and qPCR amplicons (horizontal green lines) used in the study. Note the overlap of sno-lncRNAs and *SPA2-lncRNA*. A diagram depicting the experimental design is shown on right. (B) Fold change relative to negative control (red line) in SPA- and sno-lncRNAs levels 24h post introduction of GapmeRs into iPSCs. qPCR analysis showing an average of three independent experiments, error bars correspond to standard error. (C) chRNA-seq reads from SPA-sno-lncRNAs host gene *SNHG14* in iPSCs treated with different sets of GapmeRs. Rep –replicate. chRNA-seq tracks shows counts ×10^6^.

We nucleofected iPSCs with each set of GapmeRs and isolated total and chromatin-associated fractions after 24 hours. RT-qPCR performed on total RNA revealed efficient knockdown of all seven lncRNA species using GapmeRs set 1 (SPA/sno-lncRNAs KD) (Figure 2B). However, in cells transfected with set 2 (sno-lncRNAs KD) both sno-lncRNAs and *SPA2-lncRNA* were knocked down. Set 3 (SPA-lncRNAs KD) knocked down *SPA1-lncRNA;* however, *SPA2-lncRNA* was only partially affected. Overall, we achieved a knockdown up to 60-80% of the selected RNA species across the different sets. We also tested the efficiency of GapmeRs by quantifying chromatin-associated *SNHG14* ncRNAs via chRNA-seq upon GapmeR nucleofection (Figure 2C). Our data revealed that GapmeRs set 1 efficiently reduced accumulation of *SPA1-lncRNA* (and putative *inSPA1*), *SPA2-lncRNA* and all 5 sno-lncRNAs. GapmeRs set 2 decreased sno-lncRNAs and *SPA2-*but not *SPA1-lncRNA*, while set 3 mainly affected *SPA1-lncRNA* and to a lesser extent *SPA2-* and sno-lncRNAs. Interestingly, GapmeRs-dependent knockdown was still detectable for some RNAs after 5 days from nucleofection (Figure S1B), demonstrating the utility of this system in longer experimental setups. Overall, we showed that GapmeRs can be used as an easy-to-employ alternative to genomic deletions in PWS studies.

### SPA- and sno-lncRNAs regulate transcription of neuronal genes

We analysed how rapid depletion of SPA- and sno-lncRNAs affected RNA levels in iPSCs. We did not detect global effects on the steady-state transcriptome in iPSCs nucleofected with GapmeRs set 3 targeting all PWS lncRNAs (Figure S2A). Previous studies revealed that genes regulating neuronal processes and genes contributing to immune response were affected in *post-mortem* hypothalamic tissue of patients with PWS (Bochukova et al., 2018). However, it is not clear what process drives this deregulation. Since many non-coding RNAs control transcription of protein-coding genes (Werner and Ruthenburg, 2015), next we tested levels of chromatin-associated RNAs upon SPA- and sno-lncRNAs knockdowns.

Our chRNA-seq performed on cells treated with GapmeRs sets for 24 hours uncovered the impact of ncRNAs on transcription of protein-coding genes (Figure 3A). We used differential gene expression analysis to compare SPA/sno-lncRNAs KD (set 1), sno-lncRNAs KD (set 2) and SPA-lncRNAs KD (set 3) with a control treated with non-specific GapmeRs, all in two biological repeats. We observed the greatest alterations in RNA levels between SPA/sno-lncRNAs KD and control conditions, with 205 downregulated and 87 upregulated transcripts. The differences observed between the remaining two conditions and control were more subtle, which is consistent with the varied efficiency of the GapmeR sets in depletion of PWS ncRNAs. Among the top ten most significantly downregulated genes in SPA/sno-lncRNAs KD were *FAT3, NRXN1, NLGN1* and *SNHG14* confirming the efficiency of the knockdown in this condition (Figure 3A-B and S2B). The three other top downregulated genes are involved in the regulation of neuronal development and function; *FAT3* is a cadherin which determines the polarity of developing neurons by regulating the interactions between neurites (Deans et al., 2011), while *NLGN1* and *NRXN1* are membrane adhesion proteins, which have the capacity to bind to each other across a synapse. *NLGN1* induces the formation of presynaptic boutons allowing for neuron maturation (Wittenmayer et al., 2009), and *NRXN1* regulates neuronal development, function and synaptic transmission (Uchigashima et al., 2020; Zhang et al., 2018). The comparison between sno-lncRNAs KD and control rendered only 2 downregulated genes: *FAT3* and *NRXN1*, while the SPA-lncRNAs KD failed to produce any significant changes in transcription (Figure 3A).

**Figure 3.**
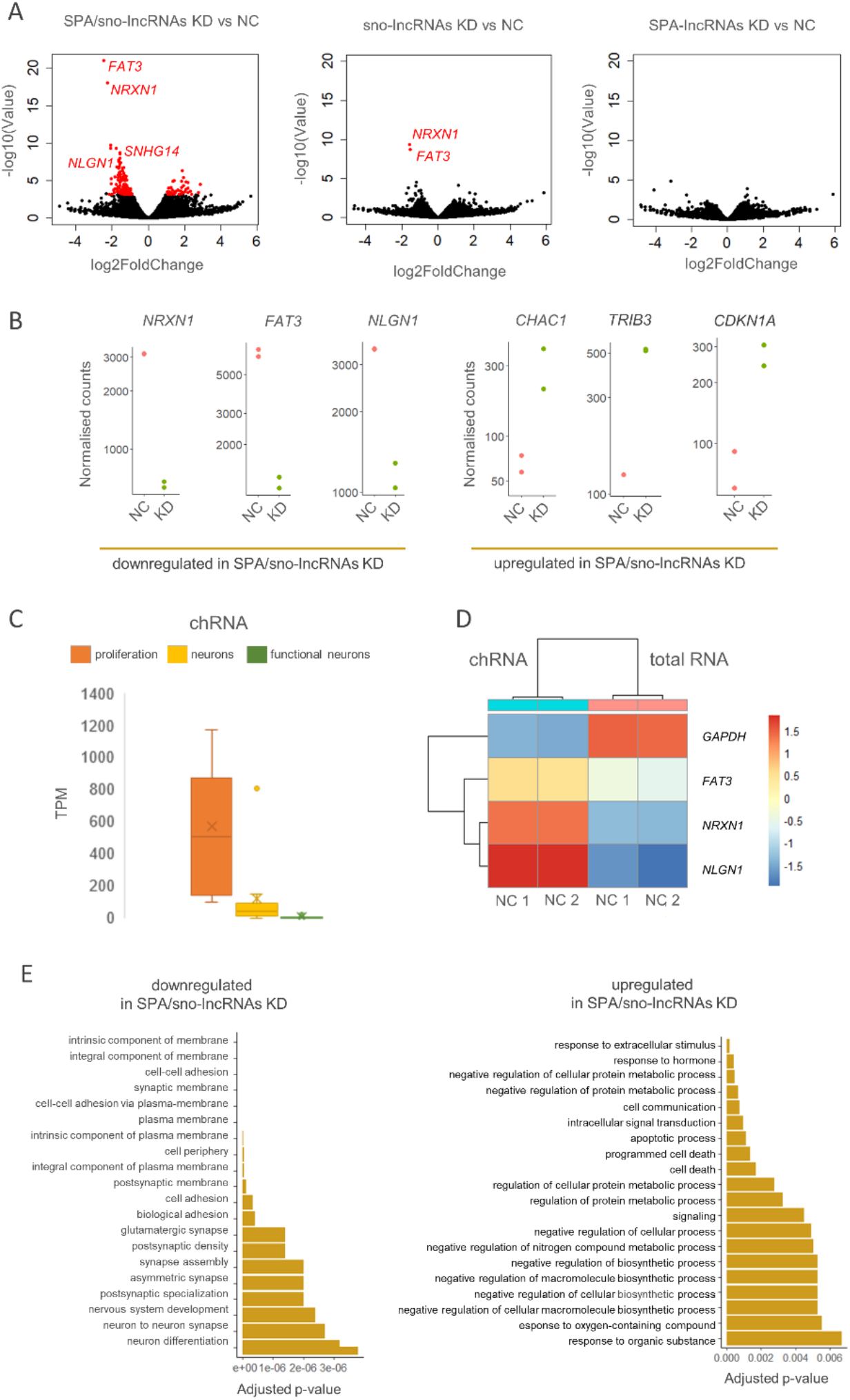
Depletion of SPA- and sno-lncRNAs affects the accumulation of chromatin-associated RNA in iPSCs. (A) Differentially accumulated chromatin-associated RNAs (in red) between the control (NC – nonspecific GapmeRs control) and SPA/sno-lncRNAs knockdown (KD) combinations. Volcano plots showing results of two biological replicates of chRNA-seq for each sample. (B) Normalised counts for genes selected from the top ten either down- or upregulated genes in chRNA-seq analysis of SPA/sno-lncRNAs depletion. (C) Transcript per million (TPM) values for markers associated with proliferation, immature, mature and functional neurons in chromatin-associated RNA fraction in iPSCs. (D) Heatmap showing the expression of neuronal and control genes in total RNA and chromatin-associated RNA fractions in iPSCs. (E) Go terms associated with downregulated and upregulated genes in chRNA-seq datasets. chRNA- chromatin-associated RNA

Interestingly, among the top ten most significant genes upregulated by SPA/sno-lncRNAs KD were genes that negatively affect proliferation and differentiation, or contribute to neuronal function: *CDKN1A, JDP2, CHAC1* and *TRIB3* (Figure 3B and S2B). *CDKN1A* negatively affects cellular proliferation by binding to and inhibiting cyclin-dependent kinases activities (Al Bitar and Gali-Muhtasib, 2019; Kreis et al., 2019). *JDP2* is involved in transcriptional responses associated with transcription factor AP-1, such as induced apoptosis and cell differentiation (Kim et al., 2010). *CHAC1* inhibits Notch signalling to promote neuronal differentiation, while *TRIB3* is an inactive kinase that plays a role in programmed neuronal cell death (Jin et al., 2002; Ohoka et al., 2005). Two other knockdowns of sno-lncRNAs and SPA-lncRNAs (set 2 and 3) did not result in transcriptional upregulation for any genes (Figure 3A). Consistent with the results of differential expression analysis, hierarchical clustering correctly grouped the duplicates of the SPA/sno-lncRNAs KD, sno-lncRNAs KD and control samples. The SPA-lncRNAs KD samples duplicates did not cluster well, which may be the result of less efficient and variable knockdowns (Figure S3A).

Active transcription of genes involved in neuronal differentiation in iPSCs was somewhat unexpected. Thus, we sought to confirm the stem status of the CREM003i-BU3C2 cell line and tested for the presence of actively transcribed neuronal markers. We calculated the abundance of transcripts in chromatin-associated RNA isolated from cells nucleofected with control GapmeRs. Transcription of proliferation markers including *POU5F1, NANOG*, and *SFRP2* was evident in the chRNA-seq data (Figure 3C and Table S1). Markers of both immature neurons (e.g. *NEUROD1, NCAM1, DCX*) and mature neurons (e.g. *ENO2, MAP2 TUBB3, NEFL*) were detected to a lesser extent in the chromatin-associated RNA fraction, and markers for functional neurons (e.g. *CHAT, TH, GAD2*) were virtually absent. A similar trend was observed in the total RNA fraction (Figure S3B and Table S1). Moreover, the accumulation of mRNA of genes involved in neurodevelopment *NRXN1, NLGN1* and *FAT3* in total RNA fraction was significantly lower than in the chromatin-associated RNA samples, which was the opposite pattern observed for constitutively expressed gene *GAPDH* (Figure 3D). This indicates that genes involved in neurodevelopment were transcribed but their mRNAs did not accumulate in non-differentiating iPSCs.

We observed the most significant differences in chromatin-associated RNA upon combined SPA/sno-lncRNAs depletion thus, we focused on exploring the functions of differentially expressed genes in this condition only. We employed Gene Ontology (GO) terms analysis to identify the relevant pathways. The genes whose transcription was downregulated by the SPA- and sno-lncRNAs knockdown are involved in pathways related to neuronal development, function and cell adhesion (Figure 3E), which is consistent with the decreased levels of *FAT1, NRXN1* and *NLGN1* in this condition (Figure 3B). These pathways include maintenance of membrane integrity, glutamatergic synapses (P < 0.001), synaptic membranes (P < 0.001), and synapse assembly processes (P < 0.001). Genes which were upregulated by the SPA- and sno-lncRNAs knockdown participate in the dampening of cellular growth and proliferation as well as in promoting apoptosis, including regulation of cellular protein metabolic processes, apoptotic processes, and programmed cell death (P < 0.001). This is consistent with transcriptional upregulation of *CDKN1A, TRIB3* and *JDP2* in this knockdown (Figure 3B).

To complement the GO analysis, we employed Gene Set Enrichment Analysis. The input for this analysis consisted of all detected transcripts, which allowed for identifying small, coordinated changes that would not otherwise be recognised (Subramanian et al., 2005). These transcripts were then ranked based on the log_2_ fold change which allowed for the determination of a wide range of activated or suppressed pathways (Figure 4A). The results of this analysis were consistent with the GO results, revealing pathways associated with central nervous system development (P < 0.001) were among those supressed upon SPA/sno-lncRNAs knockdown. This method allowed us to identify a number of other affected pathways, which were not evident from GO analysis. We found that SPA- and sno-lncRNAs depletion also suppresses pathways associated with intracellular and cell-cell signalling (correlated with terms “enzyme linked receptor protein signalling pathway” and “endosome”), as well as processes linked to immune processes (terms “leukocyte activation”, “immune effector process”), which is consistent with the misregulation of immune system genes observed in PWS (Bochukova et al., 2018). We further identified a group of activated pathways that are involved in hormonal regulation. However, these pathways included fewer genes and were associated with higher p-values than supressed pathways. Genes identified in the top five most significant pathways were visualised as cnetplot (Figure 4B). Interestingly, the genes contributing to the suppressed pathways that regulate the development of the central nervous system included *SHANK2* and *CNTNAP2*, deletions of which are associated with autism and intellectual disability (Berkel et al., 2010; Canali et al., 2018). Overall, our analyses indicate that the lack of ncRNAs transcribed from the locus deleted in PWS deregulates a broad spectrum of pathways that may contribute to aberrant development of the structures in the human brain.

**Figure 4.**
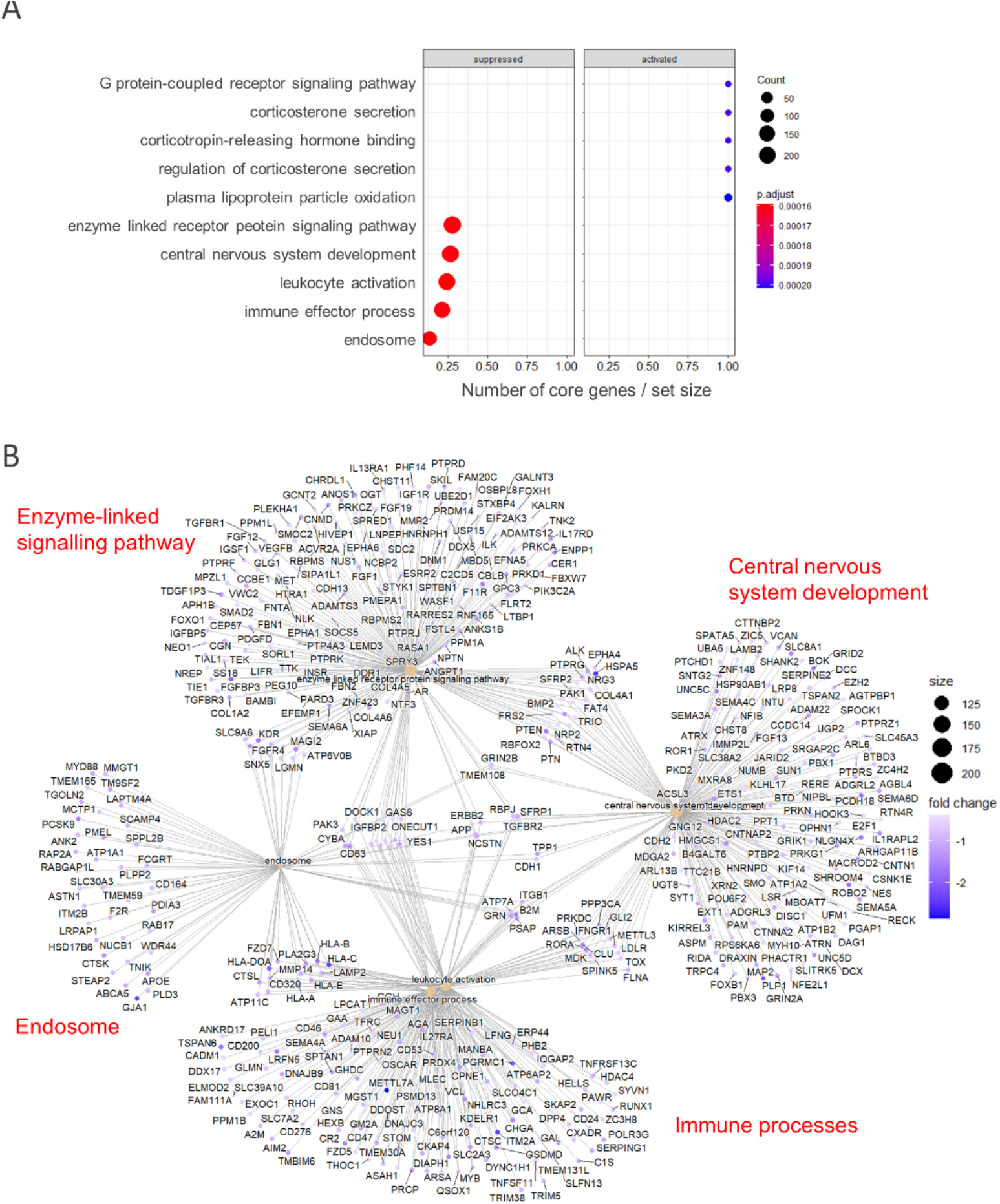
SPA- and sno-lncRNAs impact multiple pathways in iPSCs. (A) Activated and suppressed pathways indicated by changes in chromatin-associated RNA fractions. (B) Cnetplot showing genes participating in the top five supressed pathways by SPA- and sno-lncRNAs knockdown.

## DISCUSSION

The molecular basis of the neurodevelopmental genetic disorder Prader-Willi syndrome remains largely unknown. We depleted SPA-lncRNAs and sno-lncRNAs transcribed from the 15q11-q13 locus deleted in PWS to study their impact on transcription in human iPSCs. Our analysis of chromatin-associated RNA shows that the lack of these ncRNAs decreased transcription of neurodevelopmental genes and increased transcription of factors that negatively affect cellular growth and mediate apoptosis. The region downstream of the *SNURF-SNRPN* gene, *SNHG14*, that encompasses the minimal deletion resulting in PWS (Sahoo et al., 2008), produces one of the most abundant chromatin-associated RNAs in iPSCs (Figure 1). This is consistent with previous reports showing that SPA- and sno-lncRNAs levels are among the most highly expressed lncRNAs in embryonic H9 cells (Wu et al., 2016; Yin et al., 2012). The chromatin association of PWS ncRNAs points towards their possible function in transcription regulation; approximately 60% of lncRNAs display a strong bias for the chromatin association and play roles in the activation of neighbouring or distant genes (Werner and Ruthenburg, 2015).

SPA- and sno-lncRNAs display unique structural properties: the snoRNA structures at 5’ or both ends make them relatively resistant to prevalent exonucleolytic activities. Thus the intervening sequence can be used as a sponge RNA to sequester transcription and splicing factors TDP43, RBFOX2, hnRNP M and hence affect gene expression via regulation of alternative splicing (Wu et al., 2016; Yin et al., 2012). Our analysis of chromatin-associated RNA, that can be used as a proxy for active transcription of protein-coding genes (Mayer et al., 2015), uncovered that SPA- and sno-lncRNAs control the transcription of many genes that regulate neurodevelopment including *NRXN1, NLGN1*, and *FAT3*. We found that SPA- and sno-lncRNAs depletion decreased transcription levels for many genes that contribute to the formation of neuron specific structures – axons and synapses, as well as genes required for proper cell adhesion and cell-cell signalling (Figure 3 and 4). Interestingly, a significant population of genes which were upregulated by SPA- and sno-lncRNAs knockdown, are involved in the negative regulation of cellular metabolism and apoptosis. One possible explanation of this phenotype is that misregulation of neurodevelopmental genes that may result in abnormal differentiation is countered by repression of cell growth and ultimately, cell death. Indeed, neurons developed from iPSCs that model neurodevelopmental diseases including Fredreich ataxia and spinal muscular atrophy are more prone to senescence and apoptosis (Igoillo-Esteve et al., 2015; Ohashi et al., 2018). We cannot exclude the possibility that this regulation is directly provided by the factors that bind to SPA- and sno-lncRNAs however, it is more plausible that these proteins control alternative splicing of other transcriptional factors governing transcription of neuronal genes.

Interestingly, genes involved in neuron maturation and maintenance such as *NRXN1, NLGN1*, or *FAT3* were detected in the chromatin-associated fraction, indicating their active transcription (Figure 3A-B). However, their presence was not reflected at similar levels in the steady-state RNA population (Figure 3D). The accumulation of cytoplasmic RNAs may be buffered by degradation pathways that downregulate excessive transcription (Singh et al., 2019). Keeping genes that contribute to cell differentiation in the “on” state may facilitate quick transitions during development. We speculate that the transcriptional activation of genes required for neuronal development primes their efficient expression when necessary.

The global effect of SPA- and sno-lncRNAs depletion on RNA levels was completely lost in the total RNA fraction, representing mainly steady-state cytoplasmic RNAs (Figure S2A). This is consistent with the observation in a mouse model where only seven genes including transcription factor *Mafa* and growth suppressor *Necdin* were upregulated by deletions in PWS locus (Zahova et al., 2021). In human cells, deregulation of transcription caused by the absence of SPA- and sno-lncRNAs may feature in total RNA levels in later stages of neurodevelopment, when the expression of these genes is essential to support neuronal maturation. Moreover, adjusting mRNA to optimal concentration in dynamically differentiating stem cells, when the gene transcription is affected, may not be as responsive and efficient as in healthy cells. Thus, it may introduce errors in gene expression that accumulate during development and, as a consequence, manifest as PWS. Such a pathological pattern, where a transcriptional regulation imbalance emerges already in stem cells, impacts neuronal development and results in a late onset disease, has been reported for many neurodevelopmental disorders including Fragile X and Rett syndromes as well as neurodegenerative Alzheimer’s and Huntington’s diseases (Sabitha et al., 2021; Sorek et al., 2021). Mild but persistent effects on overall gene expression may be the reason why the deletions in the PWS locus are not lethal. However, as neurons progress through highly organised processes of differentiation, migration and functional activation, any disruption in gene expression may affect the development of human brain, leading to the manifestation of neurodevelopmental disorders.

## ACKNOWLEDGMENTS

This work was funded by a grant from Foundation for Prader-Willi Research (FPWR). M.S. was supported by FPWR, DH is funded by EPSRC (grant EP/T002794/1), BBSRC (grants BB/L006340/1 and BB/M017982/1); MJ is supported by Midlands Doctoral Training Partnership for a studentship to M.J. funded by BBSRC BB/M01116X/1. R.A. was funded by the Royal Embassy of Saudi Arabia Cultural Bureau and Saudi Arabia Ministry of Higher Education. P.Grz. was supported by a Sir Henry Dale Fellowship from the Wellcome Trust and the Royal Society (200473/Z/16/Z). We thank George Murphy for sharing the iPSC line. We thank Kinga Kamieniarz-Gdula and Holly Fagarasan for critical reading of the manuscript and Genomics Birmingham at the University of Birmingham for assistance with NGS.

## AUTHOR CONTRIBUTION

M.S. performed experimental part of this study. M.S., M.J. and D.H. analysed the sequencing data. R.H. and P.G. assisted with iPSCs culturing. M.S., P.G. and P.Grz. design the experiments. M.S. and P.Grz. wrote the manuscript. P.Grz. conceived the project.

**Figure S1.**
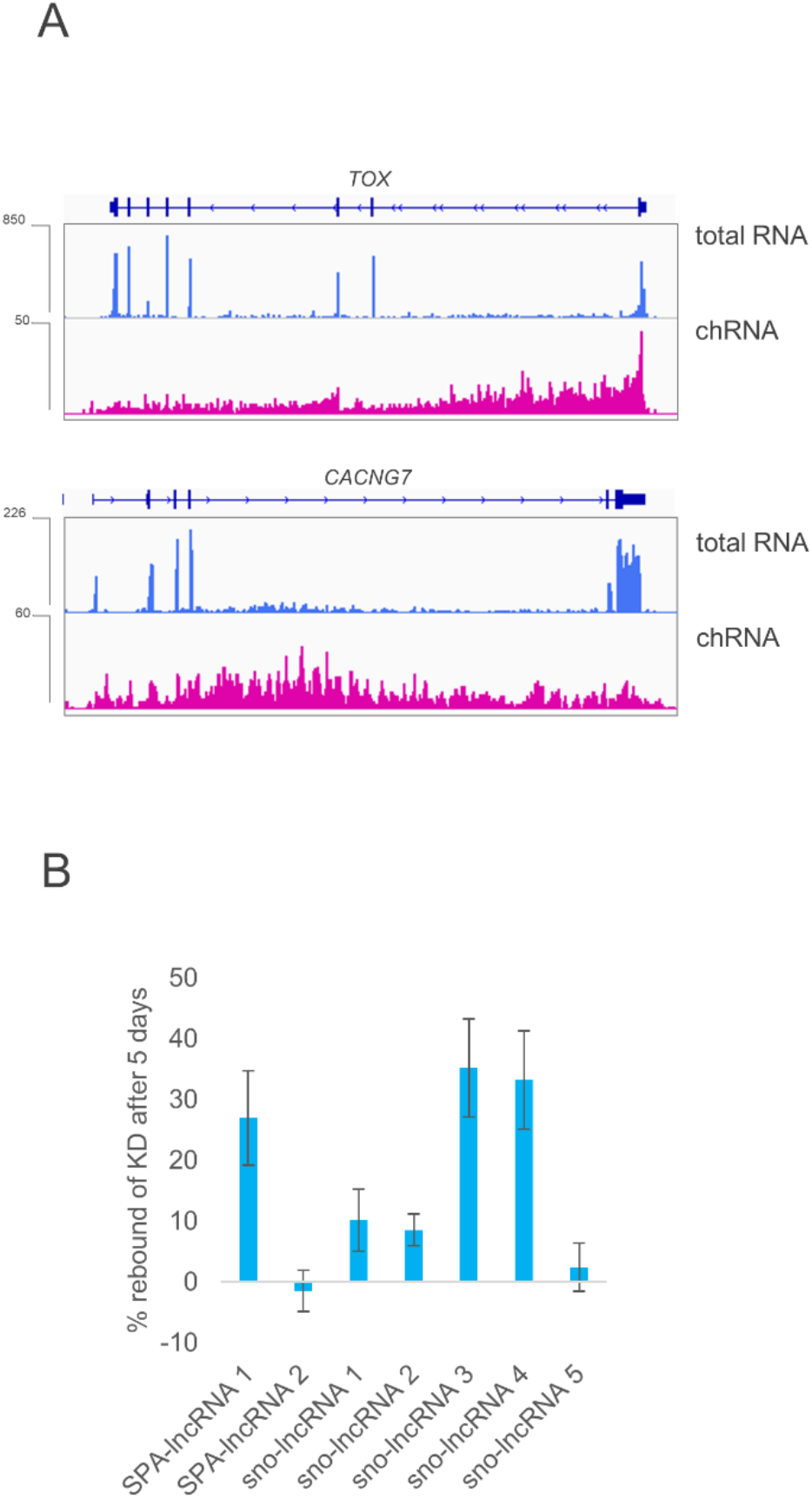
(A) Reads for *TOX* and *CACNG7* in total and chromatin-associated RNA (chRNA) samples showing enrichment of reads in the intronic regions, characteristic for samples containing nascent RNA. chRNA-seq analysis. (B) Percentage increase in expression of sno/SPA-lncRNAs between 24 h and 5 days post knockdown. RT-qPCR analysis. chRNA-seq track shows counts ×10^6^.

**Figure S2.**
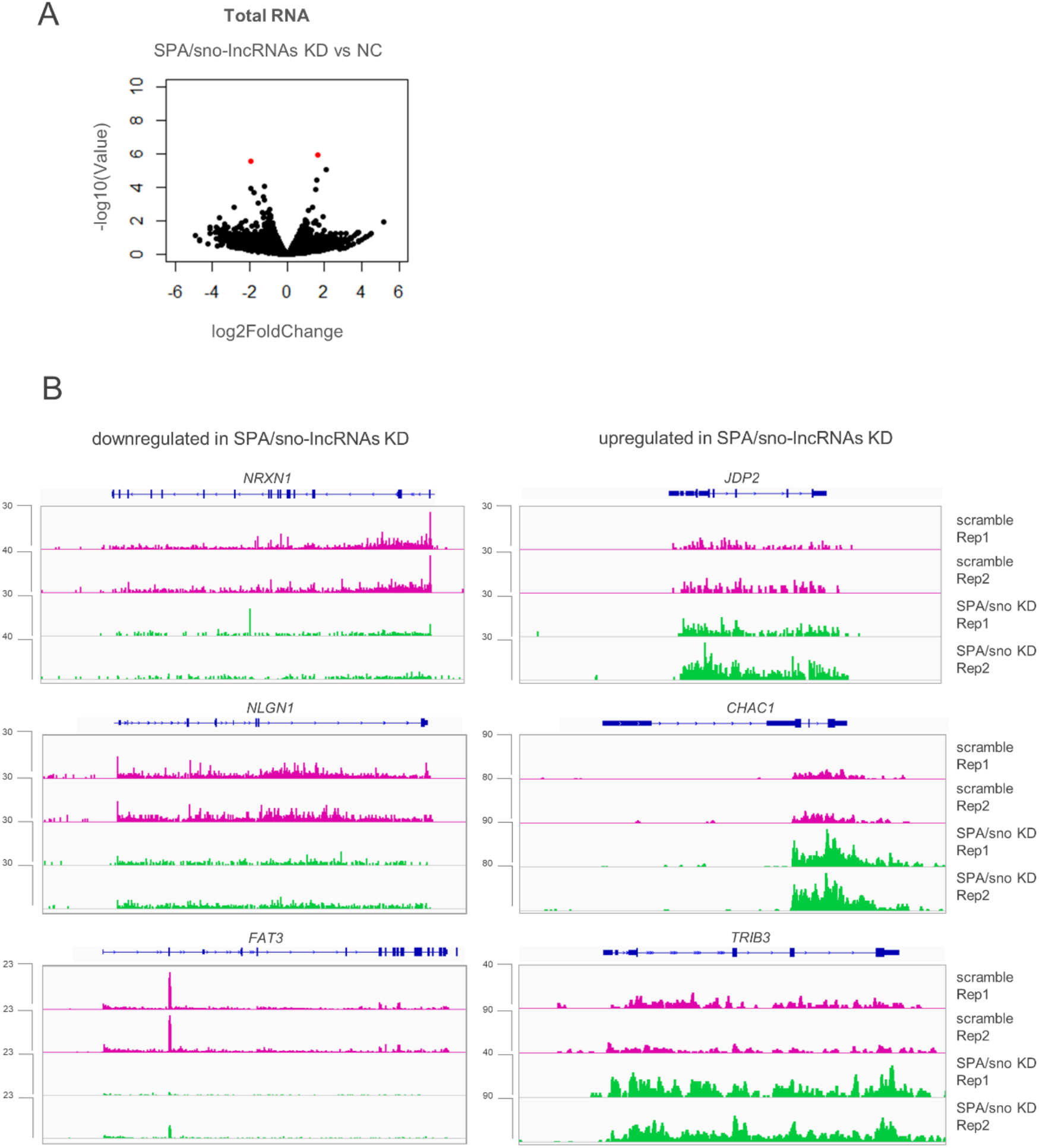
(A) Volcano plot showing differentially expressed genes (red) between control and SPA/sno-lncRNAs knockdown in total RNA fraction. (B) chRNA-seq reads for the top downregulated genes in SPA/sno-lncRNAs KD and control. (C) chRNA-seq reads for the top upregulated genes in SPA/sno-lncRNAs KD and control. chRNA-seq track shows counts ×10^6^.

**Figure S3.**
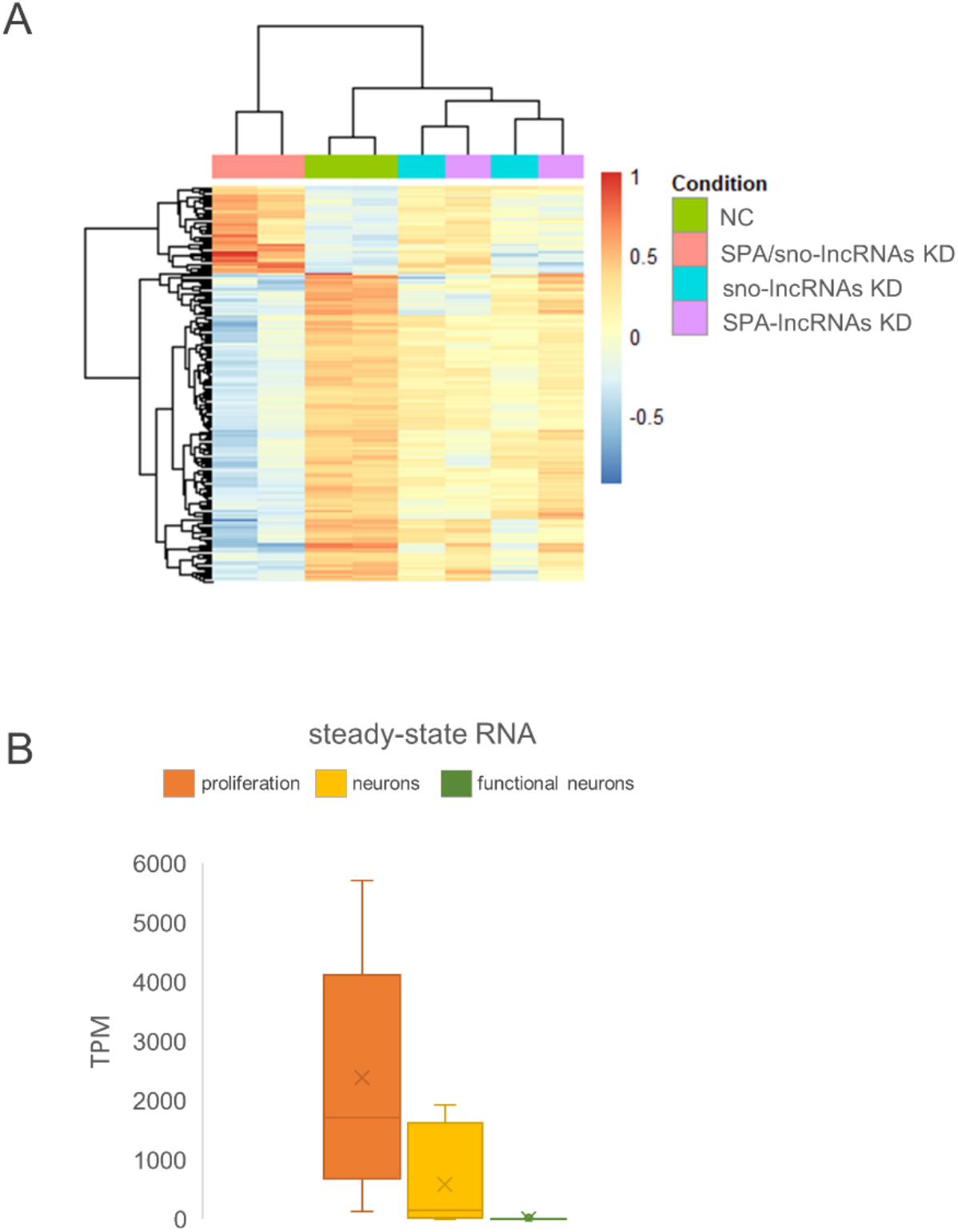
(A) Heatmap of all of the differentially expressed genes between sno/SPA-lncRNAs KD and control in chRNA-seq. Note correct clustering between the sno/SPA-lncRNAs and control samples. (B) Transcript per million (TPM) values for markers associated with proliferation, immature, mature and functional neurons in control total RNA samples.

**Table S1.** Proliferation, neurons and functional neurons markers.

| **Proliferation** | **Neurons** | **Functional neurons** |
| --- | --- | --- |
| POU5F1 | NEUROD1 | CHAT |
| NANOG | NCAM1 | TH |
| CNMD | DCX | GAD2 |
| USP44 | ENO2 | SLC17A7 |
| VSNL1 | MAP2 | SLC17A6 |
| CRABP1 | TUBB3 | FEV |
|  | NEFL |  |
|  | NEFM |  |
|  | NEFH |  |
|  | GAP43 |  |

## METHODS

### RESOURCE AVAILABILITY

#### Lead contact

Further information and requests for resources and reagents should be directed to and will be fulfilled by the lead contact, Pawel Grzechnik.

#### Materials availability

This study did not generate new unique reagents.

#### Data and code availability

The dataset generated during this study are available at GEO (GSE174043).

### EXPERIMENTAL MODEL AND SUBJECT DETAILS

Human iPSC line CREM003i-BU3C2 (Park et al., 2017) originating from a blood samples of a 40 year old human male, was kept at 36°C and 0.5% CO_2_. The cells were cultured in 6-well plates coated with Matrigel (Corning), with StemFlex medium (Gibco) supplemented with Primocin (InvovGen). The media was changed 48 h after a passage and then every 24 h. Cells were passaged using STEMPRO EZPassage tool (ThermoFisher Scientific) when 70-80% confluent. HEK293T and HeLa transformed cell lines derived from human embryonic kidney and cervical cancer cells respectively. Both were cultured at 36°C and 0.5% CO_2_, in 10cm plates, with DMEM medium (Gibco) supplemented with 10% FBS and Penicillin-Streptomycin antibiotics mixture. Media was changed every 3-4 days.

### METHOD DETAILS

#### GapmeR design

Antisense oligonucleotides (GapmeRs) were designed using QIAGEN online tool with intervening sequences of SPA- and sno-lncRNAs as input. GapmeRs for each ncRNA were selected based on the QIAGEN’s design score and they were synthesised by QIAGEN.

#### Gapmer-mediated knockdown

Confluent iPSCs were treated with 1 ml of TrypLE (Gibco) per well in 6-well plates in order to obtain a single-cell suspension. Cells were incubated for 2 min at 36°C when TrypLE was removed, and then cells were placed back at 36°C for another 3 min. 1 ml of DPBS (Gibco) was added to each well and the cells were collected and centrifuged at 200 *g* for 5 min. The supernatant was removed and the cells were resuspended in 100 µL of nucleofector solution from P3 Primary Cell 4D-Nucleofector Kit (Lonza) per transfection. The cells were then divided into individual nucleofection cuvettes, and the appropriate mixture of GapmeRs were added, 6 uL of each. Approximately 2 × 10^6^ cells were utilised per transfection. The transfections were performed using 4D-Nucleofector (Lonza) using setting DS-150. Following the nucleofection the cuvettes were incubated at 36°C for 5 min, then transferred to 1 µL of StemFlex medium and incubated at 36°C for another 10 min. The cells were plated on a Matrigel-coated 6-well plate, cells from every nucleofection were equally divided between the 6 wells in a plate. The cells were then cultured using standard conditions detailed above for 24 h after which they were collected for RNA extraction.

#### RNA extraction and fractionation

The cells were with 1 ml of TrypLE (Gibco) per well in a 6-well plates in order to obtain a single-cell suspension. The cells were incubated for 5 min at 36°C or until they detached, at which point 2 mL of DPBS was added per well. The cells were collected and centrifuged at 200 *g* for 5 min. The supernatant was removed and the cells were either resuspended in 1 mL of TRIZOL (ThermoFisher Scientific) for total RNA extraction or in 200 µL of Cytoplasmic Lysis Buffer (0.15% NP-40, 10 mM Tris-HCl pH 7, 150 mM NaCl, 50 U RiboLock) for chRNA fractionation.

For chRNA fractionation, samples were incubated for 5 min on ice in the Cytoplasmic Lysis Buffer, after which they were layered on 500 µL of Sucrose Buffer (10 mM Tris-HCl pH 7, 150 mM NaCl, 25% Sucrose, 50 U RiboLock). Nuclei were collected by centrifugation of 16000 *g* for 10 min at 4°C. The supernatant containing cytoplasmic fraction was then removed, and nuclei were washed with Nuclei Wash Buffer (PBS supplemented with 0.1% Triton X-100, 1 mM EDTA, 50 U RiboLock) at 1200 *g* for 1 min at 4°C. The supernatant was discarded and the nuclei were resuspended in 200 µL of Glycerol Buffer (20 mM Tris-HCl pH 8, 75 mM NaCl, 0.5 mM EDTA, 50% glycerol, 0.85 mM DTT, 50 U RiboLock). Next 200 µL of Nuclei Lysis Buffer (1% NP-40, 20 mM HEPES pH 7.5, 300 mM NaCl, 1 M Urea, 0.2 mM EDTA, 1 mM DTT, 50 U RiboLock) was mixed with samples by pulsed vortexing for 2 min, and centrifuged at 18500 *g* for 2 min at 4°C. The pellet containing chRNA was resuspended in 200 µL of PBS supplemented with 50 U RiboLock. Following resuspension, 500 µL of TRIZOL was added and the samples were vortexed.

At this point, for both total RNA and chRNA samples, 100 µL of chloroform was added and the samples were incubated at room temperature for 5 min. The samples were then centrifuged at 16000 *g* for 15 min at 4°C. The samples were then processed using RNeasy kit (QIAGEN), following the “RNA clean-up” protocol enclosed with the kit. The samples were then quantified using NanoDrop spectrometer and gDNA contamination was removed using TURBO DNA-free Kit (ThermoFisher Scientific) according to the manufacturer’s instructions. The samples were stored at -80°C until they were further processed for RT-PCR or RNA sequencing.

#### RT-qPCR

The gDNA-depleted RNA samples were reverse transcribed using SuperScript III Reverse Transcriptase (ThermoFisher Scientific). Briefly, 500 – 2000 ng of RNA were diluted up to 11 µL of RNAse-free water, combined with 1 µL of Random Hexamer Primers (Thermofisher Scientific) and 1 µL of 10 mM dNTPs and incubated at 65 °C for 5 min. Following the incubation the samples were briefly placed on ice, and 4 µL of SuperScript III Reverse Transcriptase buffer (Thermofisher Scientific), 2 µL of DTT, 8 U of RiboLock and 1 µL of SuperScript III Reverse Transcriptase were added per sample. The samples were incubated at 25 °C for 10 min, at 50 °C for 40 min and at 82 °C for 5 min in a thermocycler.

For RT-qPCR, the cDNA samples were diluted 1:10. For each 15 µL reaction, 7.5 µL of 2X SyGreen Mix (PCRBio), 0.8 µL of each 10 µM primer, 0.9 µL of PCR grade water and 5 µL of diluted cDNA were mixed. Three reactions per sample and per set of primers were prepared and processed using RotoGene (QIAGEN) with standard cycling.

#### RNA sequencing

Prior to RNA sequencing, total RNA samples were rRNA depleted using RiboCop (Lexogen) according to manufacturer’s instructions. Samples were re-quantified using Qubit (Thermofisher Scientific), and up to 100 ng of RNA was used for library preparation using NEBNext Ultra II RNA Library Prep Kit for Illumina (NEB). The libraries were prepared according to manufacturer’s instructions, quantified and quality checked using TapeStation (Agilent). The libraries were prepared with indexing primers and pooled into 2 library preps. There were 8 chRNA libraries that were pooled together and sequenced on a single G NextSeq 500/550 (150) flow cell to obtain sequencing depth of approximately 40M reads per sample. The 4 total RNA libraries were pooled with another 8 libraries not analysed here; the 12 libraries were then loaded and sequenced on another G NextSeq 500/550 (150) flow cell resulting in a lower sequencing depth. The RNA sequencing was performed at the Genomics Facility at the University of Birmingham. Briefly, the concentrations of the libraries were determined using Qubit and the average library size was determined using TapeStation (Agilent). The libraries were then diluted to 1.6 pM, and 1% of 20 pM PhiX control were added. The libraries were then loaded onto a flow cell and the sequencing was performed on Illumina NEXTseq.

#### Quantification and statistical analysis

Data quality control was performed with FastQC v0.11.5 (Andrews, 2010), and aligned with STAR v2.5.3a (Dobin et al., 2013) to the human genome (GRCh38.p10, Gencode comprehensive annotation). RNA-seq reads were aligned with parameters; --outSAMtype BAM SortedByCoordinate. The counts per gene were calculated with LiBiNorm v2.4 (Dyer et al., 2019), in HTseq-count (Anders et al., 2015) compatible mode with parameters; --htseq-compatible --order pos, --stranded no, --type gene, --idattr gene_name. Count per million normalised bigwig files were constructed using deeptools (Ramírez et al., 2016) bamcompare with parameters; --binSize 15, --normalizeUsing CPM. Once raw counts were extracted the data was analysed in R (version 4.0.5). Differential expression analysis was performed using DESeq2 package. A gene was considered differentially expressed when the adjusted p-value was smaller than 0.1 which is a standard setting for this package. The set of differentially expressed genes was then used as input for the GO analysis, which was executed using topGO package. All of the detected genes were used as input for GSEA analysis which was done using clusterProfiler package.

